# Courtship choreography is stabilised among genetically isolated populations

**DOI:** 10.1101/2025.09.08.675024

**Authors:** Nathan J. Butterworth, Thomas E. White, Blake M. Dawson, Jesse Appleton, Clayton McDonald, Angela McGaughran, Gregory Markowsky, Keith M. Bayless

## Abstract

Sexual selection has sculpted diverse and intricate courtship displays throughout the animal kingdom, where failure to achieve the choreographic standards of a potential partner can be highly costly for reproductive success. Yet this raises a paradox: if there is such strong selection for optimal display choreography within species, how do courtship displays diversify so extensively between species? To address this, we measure how the choreography of courtship changes among allopatric populations of the dancing dune fly – *Apotropina ornatipennis* Malloch (Diptera: Chloropidae) – a species in which males and females spend their days cavorting on Australia’s hot sandy beaches. Merging population genetics with detailed quantification of the courtship display we explore which elements of the display are the first to diverge between isolated populations, whether new behaviours arise rapidly, and whether sequence rearrangements occur in the modular structure of the display. We find that these tiny flies express courtship repertoires approaching the levels of visual complexity seen in birds of paradise. Yet despite clear genetic and geographic isolation, the complex choreography of courtship displays is stable among populations. In contrast to the notion that courtship behaviour should be highly evolvable and rapidly diverge among allopatric populations, our findings suggests that the complex choreography of courtship can instead act as a stabilising feature that limits divergence over short evolutionary timescales.

## Introduction

The animal kingdom abounds with an extraordinary variety of courtship displays that have evolved over millions of years to captivate the attention of receivers. Inadvertently, they have also captivated the attention of our own sensory systems (Darwin 1871; Bastock 1967; Thornhill & Alcock 1983; Eastwood 1999; Shuker & Simmons 2014; Arnold et al. 2017; Cannon 2023). From the tandem runs of courting *Electrotermes* termites trapped in amber deposits 38 million years ago (Mizumoto et al. 2024), to the graceful pirouettes and bows of feral pigeons and their relatives (Columbiformes) (Roberts 1905; Boehm 1955; Fabricius and Jansson 1963; Goodwin 1966; Frith 1977), to the iridescent abdomen-shaking and leg-waving displays of over 100 described species of peacock spiders (Salticidae: *Maratus*) (Girard et al. 2015; Schubert 2020; Girard et al. 2021; Otto and Hill 2021). These intricate and high-stakes performances – where *both* males and females interact with the opposite sex through synchronised movements (see Gwynne & Simmons 1990; Amundsen 2000; Amundsen & Forsgren 2001; Kraaijeveld et al. 2007; Clutton-Brock 2009; Edward & Chapman 2011) – are crucial to reproductive success (Vinnedge & Verrell 1998; Shamble et al. 2009; White et al. 2020).

While the macroevolutionary patterns of courtship display evolution are increasingly well documented across the animal kingdom (e.g., Spieth 1952; Goodwin 1966; Arnold 1972; Frith 1977; Kusmierski et al. 1997; Senter et al. 2014; Arnold et al. 2017; Ligon et al. 2018; Miles and Fuxjager 2018; Chen et al. 2019; Broder et al. 2021; Girard et al. 2021; Yukilevich 2021), we still have a rather limited understanding of the processes that generate this diversity over shorter timescales (West-Eberhard 1983; Ptacek 2000; Wiens 2000; Svensson & Gosden 2007; Wilkins 2013; Mendelson et al. 2014; Arnold & Houck 2016; Mitoyen et al. 2019; Schwark et al. 2022; Fuxjager et al. 2022; Sibly & Curnrow 2025). For example, if the precise choreography of courtship is so crucial to mating success and commonly under stabilising selection within species, then how are deviations from the optimal display selected during speciation? Over what timescales are novel behaviours invented, or choreographies rearranged (Arnold et al. 2017)? And ultimately, what microevolutionary processes drive one species to ‘pirouette’ (Fabricius & Jansson 1963) while its close congener ‘kangaroo hops’ (Cramp 1958)?

Studying differences in courtship displays among natural populations can provide crucial insights into the processes that shape their expression at the earliest branches of evolutionary divergence (Endler 1977; Foster and Endler 1999; Svensson & Gosden 2007; Mendelson et al. 2014; Iglesias et al. 2018; Gallagher et al. 2022). Empirical data shows ample evidence of divergence in courtship displays among populations (Luyten & Liley 1985; Kanmiya et al. 1990; Paillette et al. 1997; Uy and Borgia 2000; Smith & Hunter 2005; Etges et al. 2006; Arbuthnott et al. 2010; Iglesias et al. 2018; Gallagher et al. 2022) – suggesting that they can diversify over short evolutionary timescales. This high evolvability of courtship traits has been reiterated by many experimental studies (West-Eberhard 1983; Gleason & Ritchie 1998; Mendelson & Shaw 2005; Snook et al. 2005; Zuk et al. 2006; Tinghitella 2008; Rogers & Greig 2008; Arbuthnott 2009; Ding et al. 2016; Han et al. 2016; Gallagher et al. 2022) and fits neatly into the general notion that behavioural phenotypes are particularly evolutionary labile (Gleason & Ritchie 1998; Arbuthnott 2009; Blomberg et al. 2003; Hernández et al. 2021). Yet, there are also many examples where courtship displays do not diverge substantially among populations but rather appear to be consistent and stabilised (Butlin et al. 1985; Gerhardt 1991; Noor et al. 2000; Watts et al. 2019; Moran et al. 2020). This aligns with macroevolutionary evidence of courtship stability – for example, the remarkable phylogenetic conservation of the bowing display in pigeons (Columbidae) (Goodwin 1966; Frith 1977), the straddle in *Lispe* flies (Muscidae) (White et al. 2020; Butterworth et al. 2021), and the tail-straddling walk of *Plethodon* salamanders (Plethodontidae) (Arnold et al. 2017). Altogether suggesting that courtship displays (or at least some components of them) can be subject to stasis or gradual evolution.

This raises the question: When and why should we expect to see divergence of courtship displays among contemporary populations? The extent and rate of among-population divergence for any given component of a courtship display will depend firstly on many extrinsic factors. For example, current and historic geographic variation in ecological characteristics (such as brightness, background motion, colour, predation pressure, pathogens, resource availability and climate) will play a key role in shaping the extent of among-population diversification in courtship components by shifting male trait optima and female preference windows to align with the habitat (Fleishman 1988; Endler 1992; Butlin 1993; Day 2000; Yeh 2004). Specifically, as environments change across populations, the efficacy, costs, and information content of signals can shift, altering the optimal display through the eye of the receiver (i.e., sensory drive *sensu* Endler 1992; but see also Peters et al. 2007; Chung et al. 2014; Heinen-Kay et al. 2015; Morier-Genoud & Kawecki 2015; Cummings & Endler 2018; Wilson et al. 2021; Tinghitella et al. 2020; Boughman and Servedio 2022). Similarly, geographic variation in selection on environmental image detection (for detecting resources, prey, or predators) can fine tune visual preferences and sensitivity for cues that have not yet developed in the population and could facilitate the invasion of novel courtship behaviours.

Even in the absence of ecological differences among populations, courtship diversification can proceed depending on the form of selection on each component of the courtship display (i.e., stabilising, disruptive, or directional) (Gerhardt 1991; Shaw and Herlihy 2000; Kirkpatrick and Nuismer 2004; Brooks et al. 2005; Oh & Shaw 2013; Selz 2016; Arnold et al. 2017) and the shape of the preference landscape (Butlin 1993; Mendelson et al. 2014). Importantly, the form of selection may also depend on the function of a given courtship component. Display components involved in species- or mate-recognition might be under strong stabilising selection among populations, due to the potentially high costs of mating with the wrong species and hence exhibit stasis or diverge slowly (Butlin et al. 1985; McPeek et al. 2011; Wojcieszek and Simmons 2012). On the other hand, mate preference cues may be under disruptive or directional selection among populations, due to the costs of mating with locally maladapted individuals, and hence exhibit more evolutionary lability (Vortman et al. 2013; Selz et al. 2016; McClure et al. 2019). The possible rate of divergence will also be constrained by the network structure of the courtship display (Hebets et al. 2016). For example, behaviours that are connected and reinforce one another (i.e., modular components) may evolve more slowly (Hebets et al. 2016; Arnold et al. 2017) and likewise with pluripotent behaviours that serve multiple functions such as the bowing display of pigeons which are used for both social aggression and sexual courtship in certain species (Frith 1977). Components that exhibit redundancy (similar cues with the same function) and degeneracy (different cues with the same function) may show the greatest evolutionary lability among populations (Hebets et al. 2016; Hoke et al. 2019), in line with the way that gene duplications promote neofunctionalization (Arnegard et al. 2010).

There is also a major role of genetics in the diversification of courtship displays (Arbuthnott 2009; Chenoweth et al. 2010; Cande et al. 2012; Cande et al. 2014; Yeh & True 2014; Ding et al. 2016; Rossi et al. 2020; Duckhorn et al. 2022). Complex polygenic traits are expected to evolve more slowly (Orr, 2000) and evidence suggests that courtship is often polygenic (Cande et al. 2012; Rossi et al. 2020; Yeh & True 2014) *and* involves pleiotropic loci (Ding et al. 2016; Yeh & True 2014) – which may result in slow rates of divergence among populations that are hard to detect over contemporary timescales. In terms of population genetics, the extent of gene flow, drift, migration, and population size will also constrain how display traits diverge among populations (Endler 1977; Lande 1981; Thompson 1999; Sibly & Curnow 2025). With high migration and gene flow among large populations there may need to be very strong selection on courtship displays for populations to diverge in display traits (Endler 1973; Charlesworth and Lande 1982; Kirkpatrick 1996; Thompson 1999). Despite this, there is evidence of such geographic divergence in displays occurring in nature even in panmictic populations (Iglesias et al. 2018). Nevertheless, studies that correlate patterns of population structure and gene flow with courtship diversification in natural populations remain rare (Wong et al. 2004; Svensson & Gosden 2007; Iglesias et al. 2018).

Overall, we have a good understanding of what *should* shape the remarkable diversification of courtship displays that we see at macroevolutionary scales. Yet we lack fundamental data from natural populations on how the entire courtship choreography diverges. Examples of novel courtship behavioural traits or rearrangements occurring among populations and over contemporary timescales are sparse (Svensson & Gosden 2007; Svensson 2019; Gallagher et al. 2022; Gallagher et al. 2024), and few studies have considered the full scale of complexity in courtship displays (as per Lasbleiz et al. 2006; Scholes 2008a; Scholes 2008b; Arnold et al. 2017) – from qualitative differences in the presence/absence of behaviours, to quantitative changes in their frequencies and tempo, to changes in the sequence and arrangement of courtship modules. Also important to consider is that courtship involves a reciprocal back-and-forth between males and females – the behaviours of males in isolation can provide only half of the picture, yet female behaviours are rarely included in courtship ethograms (as per Lasbleiz et al. 2006). Importantly, female responses or preference windows may also diverge across populations even when male signals do not (Butlin et al. 1993; Simmons et al. 2001; Mendelson et al. 2014; Tinghitella et al. 2020; Boughman and Servedio 2022), which should also be measured to ascertain whether the function of different male behaviours across populations remains consistent (Watts et al. 2019).

Here, we use a quantitative ethological framework to describe the courtship displays of the dancing dune fly *Apotropina ornatipennis* (Malloch, 1923) (Diptera: Chloropidae), and to test how courtship divergence proceeds across natural populations. This species is commonly found along the beaches of the east coast of Australia. These habitats are particularly patchy, with a total of 10,000 beaches occupying 49.1% of Australia’s coastline, interspersed by rocky outcrops, mangroves, and other habitats (Australia State of the Environment Report, 2021). We expect that the patchy nature of these beach habitats precludes migration and gene flow among *A. ornatipennis* populations, and in turn that these allopatric populations will have rapidly evolved distinct differences in their courtship displays. By elucidating the precise mode of intraspecific diversification across the full suite of courtship components and choreography, we aim to provide insight into the mechanisms that drive the evolutionary diversification of courtship displays.

## METHODS

### Animals

*Apotropina ornatipennis* Malloch (1923) (Diptera: Chloropidae) is one of twenty-two described *Apotropina* from Australia (Ang et al. 2023). The species inhabits beaches on the eastern coast of Australia and is most abundant in sandy areas of the beach with coastal grasses (Poales). The larval life-history is entirely unknown, and the present authors have never observed any female oviposition. Adults are active year-round though at greatly reduced population densities in the winter (estimates of 1 to 2 individuals/m^2^) (pers. obs). Population size peaks in the warmer spring and summer months, with aggregations ranging from estimates of 1 to 30 individuals/m^2^ (pers. obs). Females spend their days foraging and often stand in the shade under fallen debris and leaves, while males frantically run across the hot sand and vigorously court any females they encounter (Figure 1; supplementary video). Females are usually larger than males, but both sexes exhibit pigmented wings, which are vibrated frequently by both sexes during courtship. The males will also engage in male-male combat, which appears to involve direct contact between the proboscis of both males (see supplementary video). *A. ornatipennis* is often the dominant species where it occurs – alongside terrestrial amphipods (Talitridae) and various small ants. Predators include spiders and flies of the genus *Lispe* (Diptera: Muscidae) which have been observed to opportunistically kill roaming *A. ornatipennis* males (pers. obs.). Most commonly, individuals occur in conspecific masses, with few opportunities for males to encounter other dipteran species and to misdirect courtship.

**Figure 1.**
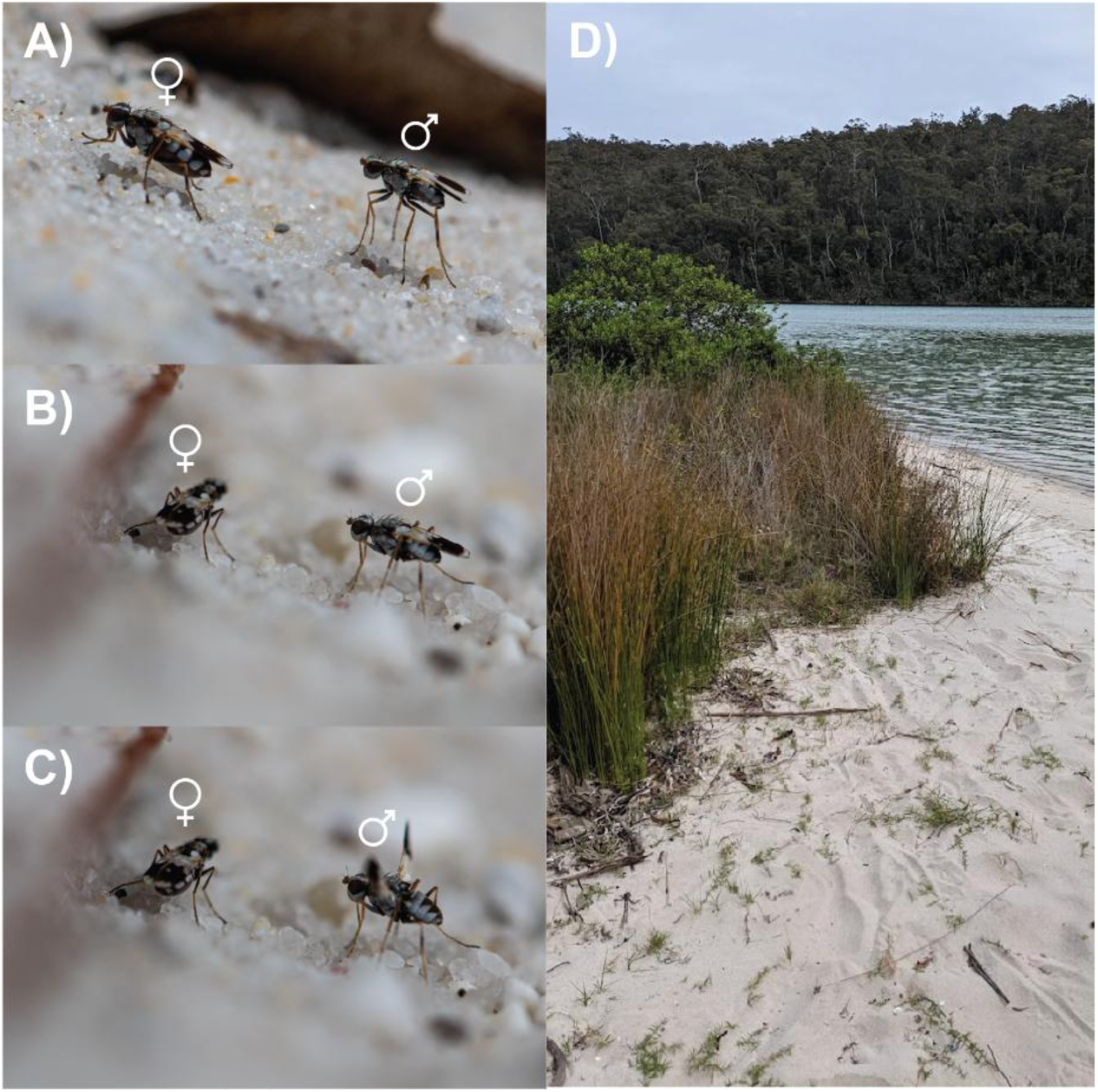
The dancing dune-fly, *Apotropina ornatipennis* (Diptera: Chloropidae). A) Orienting, B) Face-off, C) Wing-sweep, D) typical habitat.

### Population genetics

To ascertain the genetic structure of populations among beaches, flies were collected between 24^th^ of October and 17^th^ of December 2021 from seven sites (Table 1), euthanised with ethyl acetate vapour within eight hours of capture and placed into 2.5 mL plastic tubes containing 90% ethanol and stored at − 4 °C in the laboratory for up to eight weeks. A subset of specimens were taxonomically identified by comparison to the holotype and paratypes in the Australian National Insect Collection (Canberra, Australia).

**Table 1.** Summary of collection localities and population genetic parameters.

| Site | Code | Latitude | Longitude | Date collected | *N* | *H*<sub>O</sub> | *H*<sub>E</sub> | *F*<sub>IS</sub> | *F*<sub>ST</sub> | SNP<sub>PRIV</sub> |
| --- | --- | --- | --- | --- | --- | --- | --- | --- | --- | --- |
| Greenpatch | GR | -35.1365 | 150.72346 | 17/12/2021 | 23 | 0.139 | 0.19 | 0.298 | 0.0000 | 2316 |
| Wairo | WA | -35.4276 | 150.41645 | 24/10/2021 | 9 | 0.088 | 0.11 | 0.161 | 0.3547 | 122 |
| Haywards | HA | -36.4053 | 150.06543 | 24/10/2021 | 12 | 0.109 | 0.125 | 0.106 | 0.2628 | 11 |
| Beares | BE | -36.4324 | 150.07836 | 24/10/2021 | 11 | 0.106 | 0.125 | 0.135 | 0.2619 | 16 |
| Merimbula | ME | -36.9033 | 149.91 | 23/10/2021 | 19 | 0.115 | 0.131 | 0.126 | 0.2236 | 42 |
| Middle | MI | -36.8915 | 149.92931 | 12/12/2021 | 10 | 0.111 | 0.133 | 0.143 | 0.2163 | 8 |
| Severs | SE | -36.9526 | 149.90857 | 07/12/2021 | 10 | 0.108 | 0.13 | 0.158 | 0.2320 | 14 |

Each individual fly (9–23 per population) was placed in a single well of a 96-well plate with 70% ethanol, and sent to Diversity Arrays Pty Ltd (Canberra, Australia) for a high-density DArTSeq™ assay (∼ 2.5 million sequences/sample used in marker calling). The DArTSeq™ extraction and sequencing methods are detailed in Kilian et al. (2012) and Georges et al. (2018).

To ensure appropriate DNA fragments were used for subsequent sequencing, restriction enzyme digestion was optimised for *A. ornatipennis* using multiple restriction enzyme combinations and eight specimen replicates. Following sequencing of these test specimens, the optimal restriction enzyme pair was identified as PstI-HpaII, based on the fraction of the genome represented, while controlling for average read depth and the number of polymorphic loci. This restriction enzyme combination was used for all subsequent digestions. Following digestion, all sequence fragment libraries were ligated with Illumina sequencing adaptors and sequenced on an Illumina HiSeq2000 platform.

Short-read sequences were processed using the DarTseq™ bioinformatic pipeline (Georges et al. 2018), which performs filtering and variant calling, to generate final genotypes. While some parts of the sequencing and analysis protocol are proprietary and cannot be provided, the use of the DArTSeq platform for studies of genetic diversity and structure is widespread in the field (Popa-Báez et al. 2020; Hoffman et al. 2021; Jaya et al. 2022; Butterworth et al. 2023) and is reproducible.

### Genetic analysis

The DArTseq™ dataset contained a total of 38,905 SNPs across 94 individual flies (Supplementary material 1). The data were then filtered with the ‘dartR’ package version 1.9.9.1 (Gruber et al. 2018) in R version 4.5.0 (R Core Team 2025). We filtered the DArTseq™ dataset by reproducibility (proportion of technical replicate assay pairs for which the marker score was consistent) at a threshold of 0.98, then by call rate (proportion of samples for which the genotype call was not missing) at a threshold of 0.95, and finally by minor allele count at a threshold of 0.02 (MAC less than the threshold are removed). This resulted in a filtered dataset of 94 individuals, 7856 SNPs, and 1.19% missing data.

We used R for all analyses of genetic diversity. We applied the ‘basic.stats’ function of the ‘hierfstat’ package version 0.5–10 (Goudet et al. 2015) to calculate average observed heterozygosity (HO), expected heterozygosity (HE), and inbreeding coefficients (FIS). We also used the ‘betas’ function from ‘hierfstat’ to calculate population-specific FST values (Weir and Goudet 2017).

Using R, we assessed population structure by AMOVA using the function ‘poppr.amova’ with the ‘ade4’ implementation from the ‘poppr’ package version 2.9.3 (Kamvar et al. 2014). To test whether populations were significantly different, we used a randomisation test on the AMOVA output with 1000 permutations (Excoffier et al. 1992) using the function ‘randtest’ from the package ‘ade4’ version 1.7–18 (Thioulouse et al. 2018). We then conducted pairwise comparisons of FST values between populations using the ‘gl.fst.pop’ function from the ‘dartR’ package with 10,000 bootstrap replicates.

Genetic distances between individuals were examined using Nei’s distances, and a dendrogram with 1000 bootstrap replicates was created with the ‘aboot’ function of the ‘poppr’ package, and the ‘ggtree’ function of the package ‘ggtree’ (Yu 2020). We then used the ‘glPca’ function from the ‘adegenet’ package version 2.1.5 (Jombart 2008) to determine whether genetic differences between individuals (as represented by principal components) were structured according to their populations.

To test for isolation by distance, we performed a Mantel test using the function ‘gl.ibd’ from the ‘dartR’ package in R. This compared linearised genetic distances between populations (calculated using ‘StAMPP’ version 1.6.3; Pembleton et al. 2013) against Euclidean geographical distances (calculated using ‘vegan’ version 2.5–7; Oksanen et al. 2013).

To calculate individual blowfly admixture coefficients, the filtered SNP data were converted into the STRUCTURE format (‘.str’) using the ‘gl2faststructure’ function from the ‘dartR’ package, then into the ‘.geno’ format using the ‘struct2geno’ function of the ‘LEA’ package version 3.1.4 (Frichot and Francois 2015). We then ran sparse non-negative matrix factorisation on these data with the ‘sNMF’ function from ‘LEA’ to examine genetic clusters in the data. We analysed K values (i.e., cluster numbers) of 1 to 10, with 100 replications for each K value, and used the cross-entropy criterion to determine the value of K that best explained the results.

### Quantifying courtship

From the seven sites used for genetic analysis, three were chosen for detailed analysis of courtship patterns. Two adjacent populations (Severs Beach and Middle Beach) that likely exchange migrants and should therefore have relatively higher gene flow, were compared to the most geographically distant population (Greenpatch Beach), where the proportion of migrants and gene flow would be expected to be lowest (see supplementary material; Table S1). With this comparison, we would expect the Severs and Middle beach populations to exhibit most similarity in courtship displays and be highly distinct from the courtship displays observed in the Greenpatch populations.

At all beaches, filming occurred under clear conditions between the daylight hours of 0900 and 1500 and between the 1st and 17th of December 2021. Behaviour was filmed under natural light and temperature conditions with a Google Pixel 5 (12.2 MP) or iPhone 11 (12 MP) camera recording at 60 frames per second. Filming began when a male approached a female and continued until one or both flies left the area and could no longer be observed. Once video footage was obtained, slow-motion playback with Adobe Premiere Pro allowed us to describe all inter- and intra-sexual interactions (Table 2). For each male-female pair (n=15 per population) we used Solomon Coder 17.03.22 (Péter 2017) to score the durations and frequencies of all male and female behaviours during interactions. All scoring was done by a single researcher (JA). Across both males and females, we recorded 18 behaviours that occurred during the courtship display. These behaviours were not all mutually exclusive as different body parts could be used simultaneously (Table 2) – for example, males could ‘orient’ and ‘wing-vibrate’ at the same time.

**Table 2.**
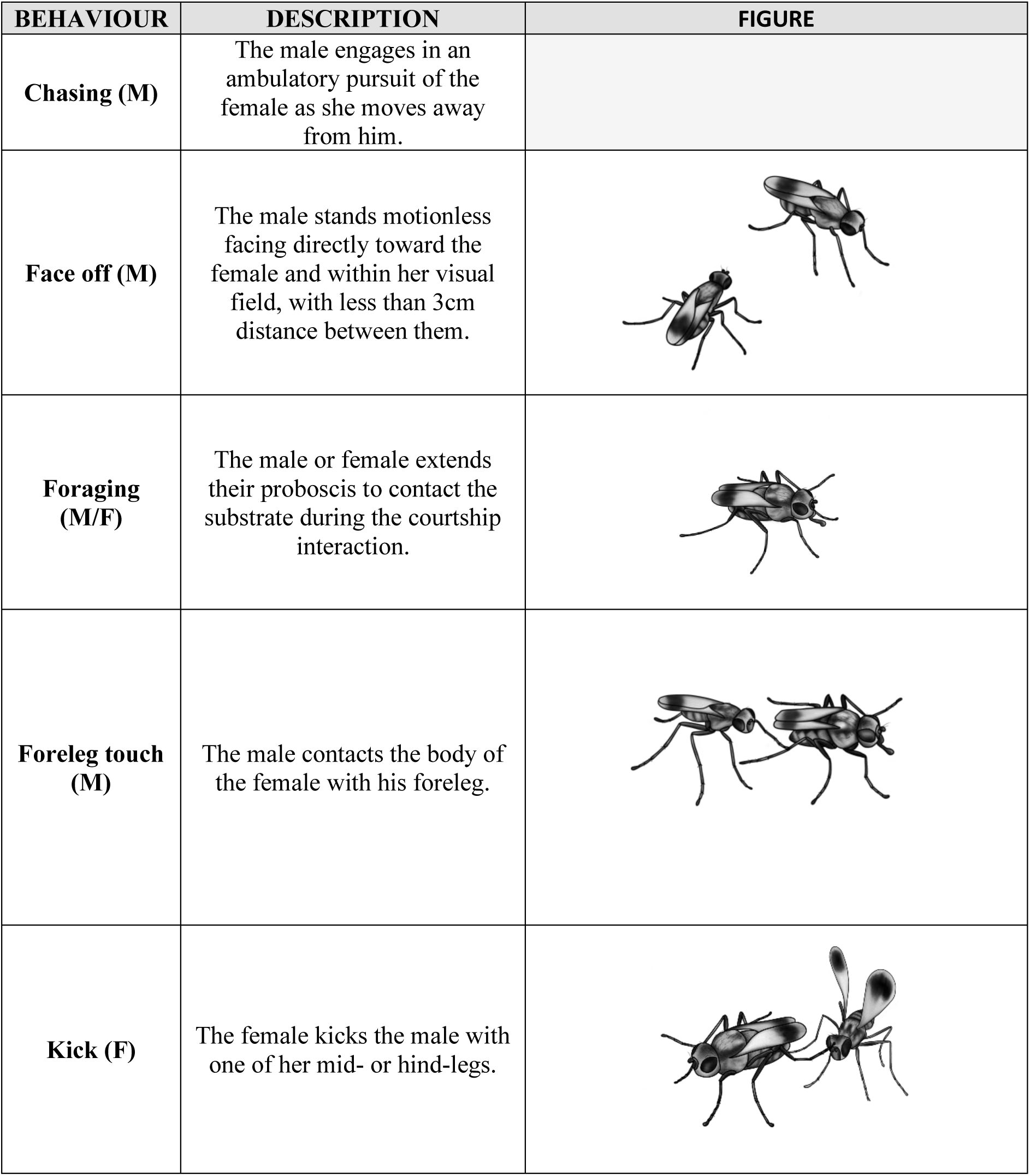
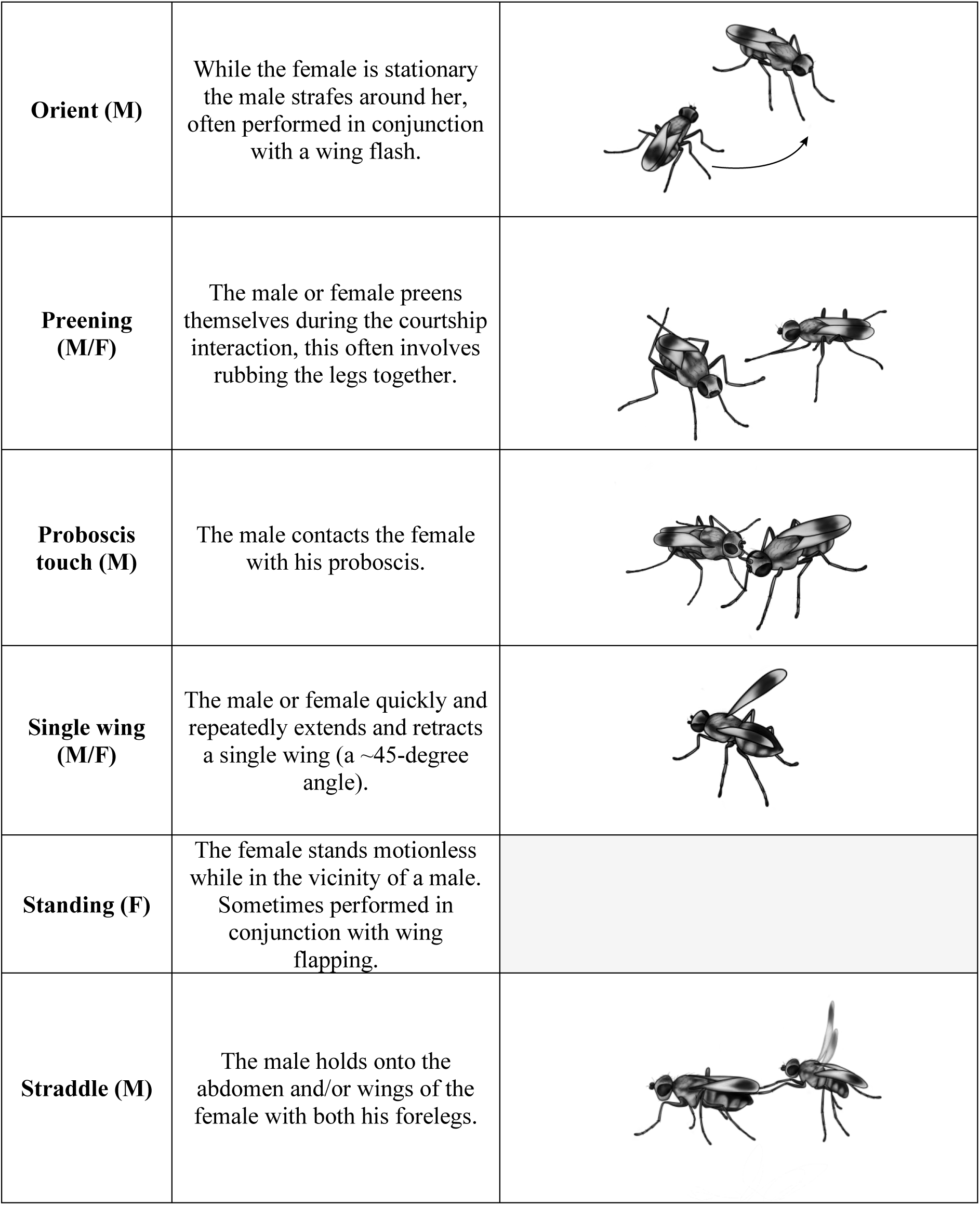
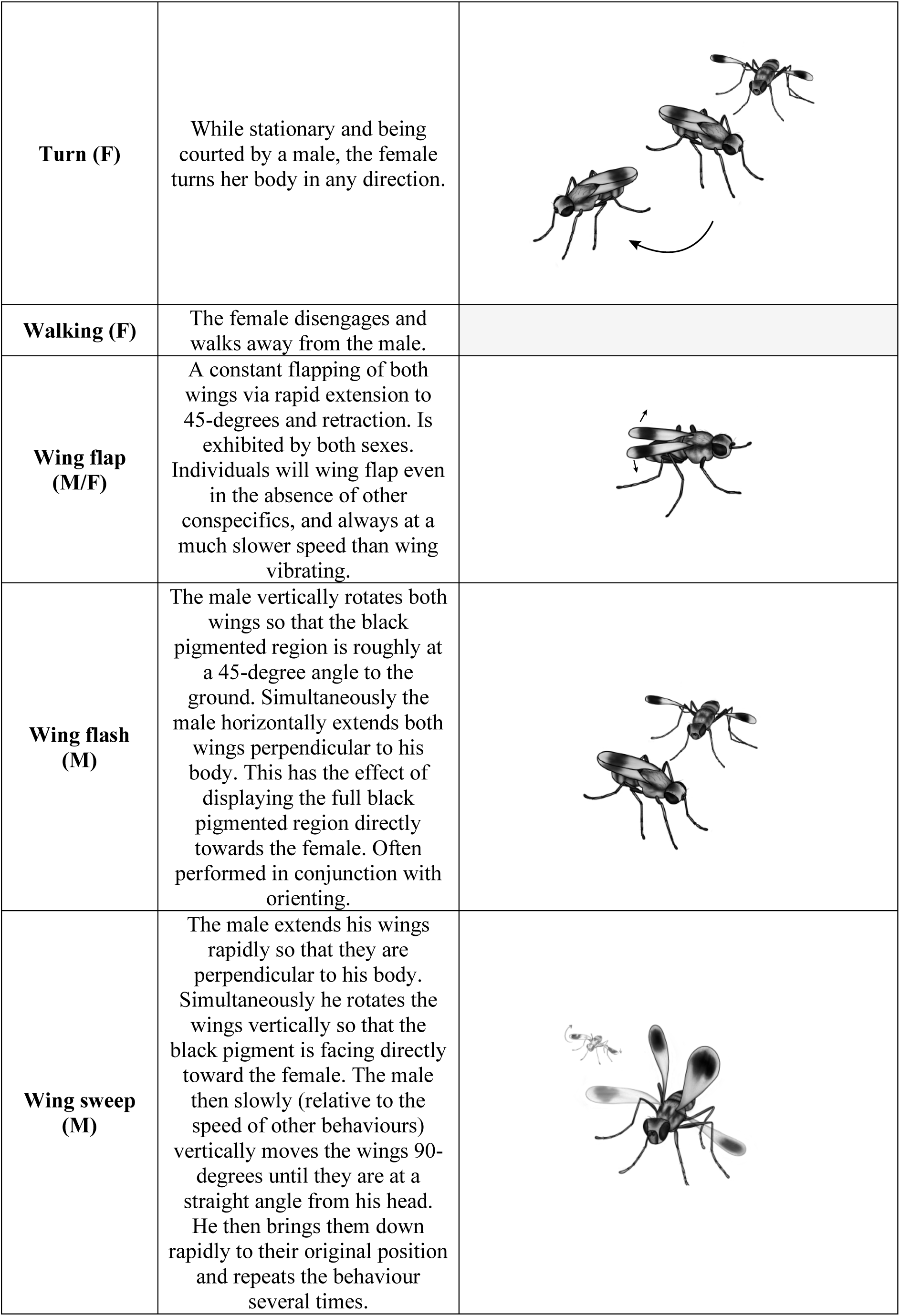
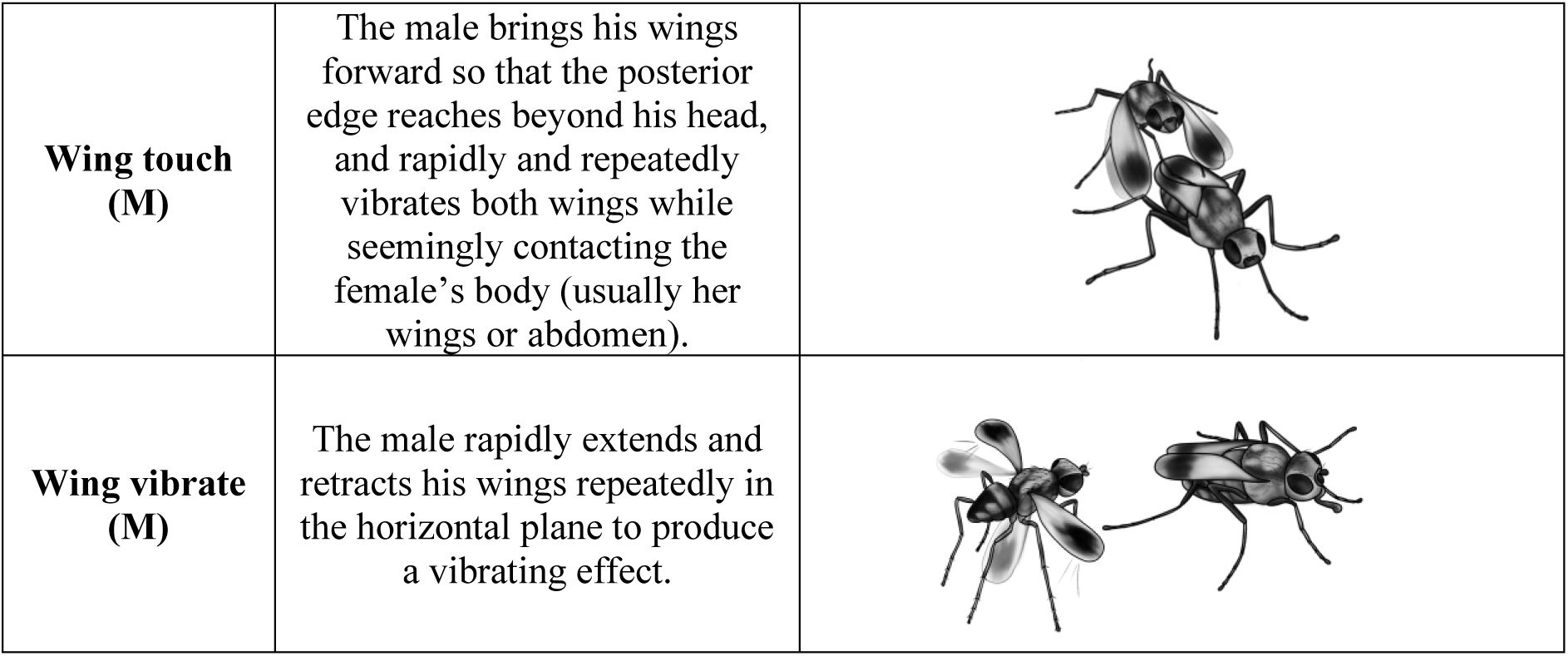
Ethogram of courtship behaviours of *Apotropina ornatipennis*. Behaviours by male (M), female (F), or both sexes (M/F).

| BEHAVIOUR | DESCRIPTION | FIGURE |
| --- | --- | --- |
| **Chasing (M)** | The male engages in an ambulatory pursuit of the female as she moves away from him. |  |
| **Face off (M)** | The male stands motionless facing directly toward the female and within her visual field, with less than 3cm distance between them. |  |
| **Foraging (M/F)** | The male or female extends their proboscis to contact the substrate during the courtship interaction. |  |
| **Foreleg touch (M)** | The male contacts the body of the female with his foreleg. |  |
| **Kick (F)** | The female kicks the male with one of her mid- or hind-legs. |  |
| **Orient (M)** | While the female is stationary the male strafes around her, often performed in conjunction with a wing flash. |  |
| **Preening (M/F)** | The male or female preens themselves during the courtship interaction, this often involves rubbing the legs together. |  |
| **Proboscis touch (M)** | The male contacts the female with his proboscis. |  |
| **Single wing (M/F)** | The male or female quickly and repeatedly extends and retracts a single wing (a ~45-degree angle). |  |
| **Standing (F)** | The female stands motionless while in the vicinity of a male. Sometimes performed in conjunction with wing flapping. |  |
| **Straddle (M)** | The male holds onto the abdomen and/or wings of the female with both his forelegs. |  |
| **Turn (F)** | While stationary and being courted by a male, the female turns her body in any direction. |  |
| **Walking (F)** | The female disengages and walks away from the male. |  |
| **Wing flap (M/F)** | A constant flapping of both wings via rapid extension to 45-degrees and retraction. Is exhibited by both sexes. Individuals will wing flap even in the absence of other conspecifics, and always at a much slower speed than wing vibrating. |  |
| **Wing flash (M)** | The male vertically rotates both wings so that the black pigmented region is roughly at a 45-degree angle to the ground. Simultaneously the male horizontally extends both wings perpendicular to his body. This has the effect of displaying the full black pigmented region directly towards the female. Often performed in conjunction with orienting. |  |
| **Wing sweep (M)** | The male extends his wings rapidly so that they are perpendicular to his body. Simultaneously he rotates the wings vertically so that the black pigment is facing directly toward the female. The male then slowly (relative to the speed of other behaviours) vertically moves the wings 90-degrees until they are at a straight angle from his head. He then brings them down rapidly to their original position and repeats the behaviour several times. |  |
| **Wing touch**<br>**(M)** | The male brings his wings forward so that the posterior edge reaches beyond his head, and rapidly and repeatedly vibrates both wings while seemingly contacting the female's body (usually her wings or abdomen). |  |
| **Wing vibrate**<br>**(M)** | The male rapidly extends and retracts his wings repeatedly in the horizontal plane to produce a vibrating effect. |  |

Over two weeks (and more than 100 total hours) of filming flies, we were only able to observe a single pair proceed all the way to copulation. This means that 44/45 of the measured courtship interactions were from unsuccessful male-female pairs. Nevertheless, trials all involved prolonged back and forth assessment between the sexes before one partner or the other left (the median trial time was 92 seconds, and the average trial time was 148 seconds) – suggesting mutual assessment and female engagement in the courtship display. All recorded courtship components have evolved under selection in these natural populations and although we may devise clearer results (particularly for female preference) if we could also assess how successful males differed in courtship among populations, this was simply not possible to achieve within the project scope. Crucially, because we observed a full courtship sequence up to mating, we can confirm that the courtship sequences we observed across all trials are the majority of what occurs during a successful display, and that mating does occur on the beach (in the same habitat as courtship interactions). Overall, we can be confident every trial analysed is an accurate representation of the courtship display and choreographic sequence of *A. ornatipennis*.

### Courtship statistical analysis

To determine whether the frequency with which behaviours occur within populations varied among populations, we qualitatively compared the proportion of flies exhibiting display behaviours among populations (number of flies that displayed behaviour /total number of flies assayed in each population).

To assess quantitative differences in the temporal patterns of male courtship displays among populations, we analysed individual behaviours (i.e., orient and wing-sweep in isolation, as opposed to the combined ‘orient-wing-sweep’) and only those that occurred above 50% frequency within a population, which allowed replication to be high enough for quantitative analysis. For each male-female courting pair we measured the following aspects of each behavioural trait: (1) the mean bout duration as a proportion of the total trial time (mean behaviour bout duration/total trial duration), (2) the frequency (number of times a behaviour occurred/total number of all behavioural occurrences), and (3) the mean inter-bout interval as a proportion of the total trial time (mean duration of time between behaviour bouts/total trial duration). All data were continuous proportions, so we used beta regression (betareg package in R; v. 3.2-4, Cribari-Neto & Zeileis 2010) to assess how each of these metrics varied among populations followed by ‘ANOVA’ type III from the ‘car’ package (v. 3.1-1, Fox and Weisberg 2019).

To assess whether female responses to male behaviours varied among populations, we compared the frequency of male wing-sweep and wing-flash against mean proportional duration per female walking bout (mean proportion of walking time per bout/total trial time) as a sign of interest. This was informed by the stationary behaviour of the single mated pair we observed where reduction in movement appeared key to mating success – the female remained stationary for 92% of the 140s trial time. We also compared the mean bout duration of male wing vibration against the mean bout duration of female wing flapping which is a common female response to courtship in flies (e.g., Butterworth et al. 2019). Because data were all in the form of continuous proportions, we used beta regression followed by ‘ANOVA’ type III, as described above.

Lastly, we assessed whether there was divergence in the structure of the courtship sequence among populations, and whether there was a relationship between genetic distance and divergence in the overall courtship sequences. Following Green and Patek (2018) we first used the igraph network analysis package (v.2.1.4, Csárdi & Nepusz 2006) to summarise behavioural sequence data into adjacency matrices for each population, where each row and column in the matrix corresponded to one of 41 behavioural combinations (41 x 41 matrix). Each cell in this matrix corresponded to the number of times, across the dataset, that one behaviour from an individual transitioned to a subsequent behaviour from that individual. We identified transitions that were more frequent than expected by chance (i.e., the non-random components of the display) using permutation procedures for sequential behavioural analysis (see Bakeman et al. 1996; Green and Patek 2018). This enabled us to isolate only the significant transitions. From these non-random transition matrices, we then used Kullback-Leibler divergences (commonly employed for comparing Markov matrices in information theory; Rached et al. 2004) to compare distances in courtship display structure between individuals within- and among-populations. We used non-informative priors for the Dirichlet distribution using ‘KL.Dirichlet()’ from the R package ‘entropy’ (v.1.3.2, Hausser et al. 2012) to calculate divergence between vectorised Markov courtship transition matrices between pairs of individuals (i.e., comparing the courtship structure of Greenpatch 1 to Greenpatch 2, then Greenpatch 1 to Severs 1, etc). To ascertain whether courtship displays divergence is related to genetic distance we then correlated these mean inter-individual courtship distances (as calculated by Kullback-Leibler divergences of transition counts per individual courting pair) with the mean inter-individual genetic distances (as calculated by Kosman distances; Kosman & Leonard 2005).

## RESULTS

### Populations are genetically isolated and exhibit low levels of gene flow

There was clear genetic differentiation among populations based on Nei’s genetic distances (Figure 2.A.i) and principal component analysis where the first two components explained 26.8% of the total variation with clear separation of populations (Figure 2.A.ii). Results of the AMOVA further indicated that populations were differentiated with 14% of genotypic variation coming from among-individual differences within populations, and 21% from among-population differences (FST = 0.2043, p = 0.001). Pairwise F_ST_ values ranged from 0.014 to 0.280 (all p-values < 0.01) and were consistently highest for comparisons involving Greenpatch populations (supplementary material; Table S1). The Mantel test indicated statistically significant correlation between genetic differentiation and geographic distance between sampled populations (p < 0.01) (Figure 2.A.iii) – though Greenpatch contrasted this overall pattern, as it was distinct even from its closest population (Wairo Beach – within 43km) and also had a high number of private alleles (Table 1). The sNMF analysis identified five genetic populations as the most likely, and Greenpatch and Wairo showed largely discrete clusters with limited admixture for K = 5 (Figure 2B). In contrast, the remaining populations showed more mixing, with shared ancestry highest at local scales (i.e., between Beares and Hayward Beach and between Merimbula, Middle, and Severs Beach).

**Figure 2.**
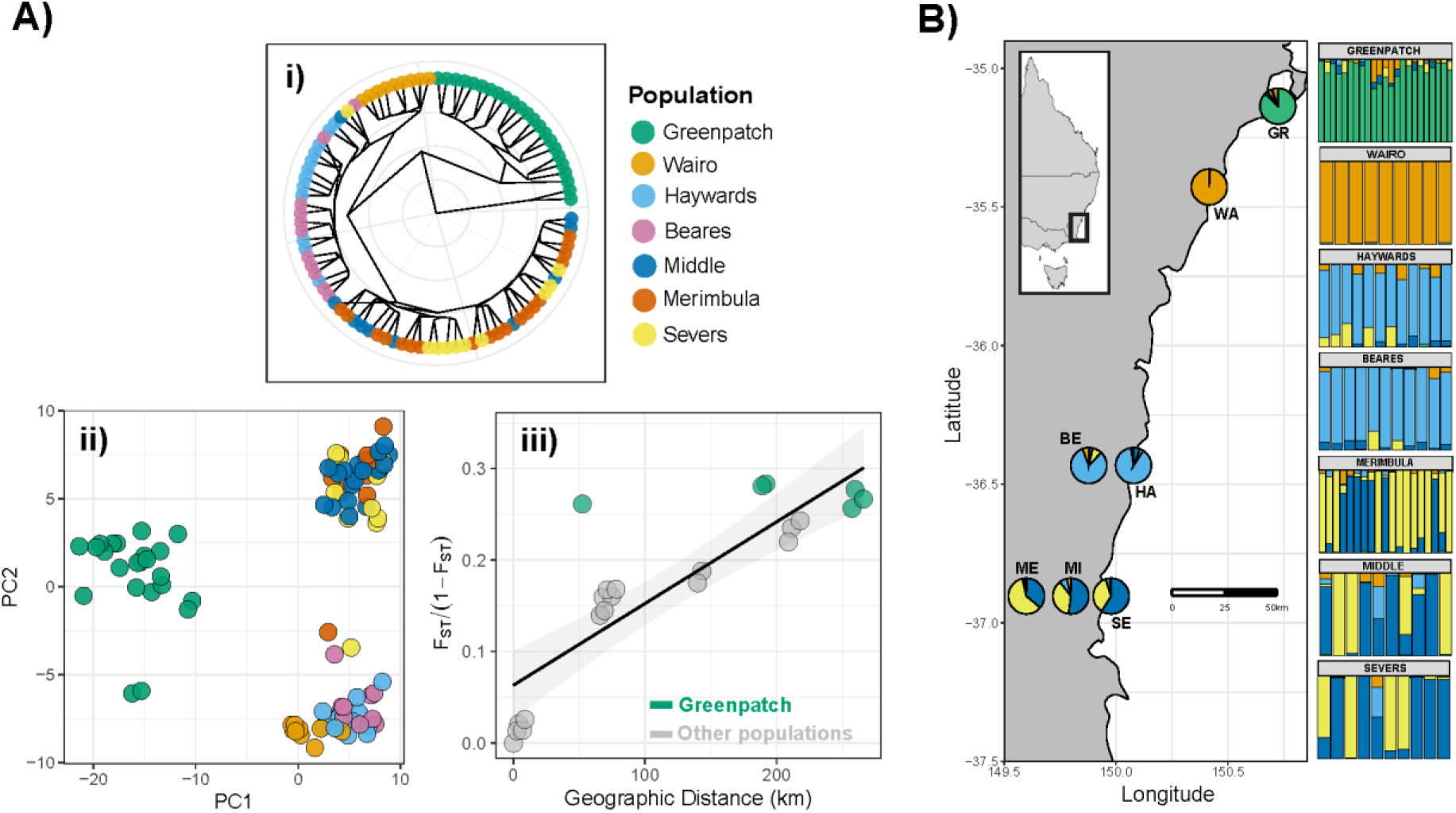
Population genetic metrics among several populations of *A. ornatipennis* (Diptera: Chloropidae). A) Genetic diversity as represented by i) Neis genetic distances, ii) Principal component analysis, and iii) A mantel test of genetic isolation by geographic distance. B) The mean admixture proportions of populations of *Apotropina ornatipennis* (Diptera: Chloropidae) that were sampled in the present study. The admixture pie charts plotted on the map represent population averages. The bar plots presented on the right reflect individual admixture proportions, sorted by population, where each bar represents a single individual. Full population names are provided in Table 1.

### The precopulatory courtship display is highly complex

Courtship was highly complex consisting of 18 discrete behaviours (Table 2), and 41 behavioural states (Table 6).

### Courtship choreography is stable among genetically isolated populations

We found no evidence of divergence among the three tested populations regarding the innovation of new behaviours, or changes in the proportion of males performing any display components. All 18 display behaviours were observed in each population at similar proportions (Figure 3). Behaviours that occurred at in more than 50% of pairs (across all sites) were used for downstream quantitative analysis among populations.

**Figure 3.**
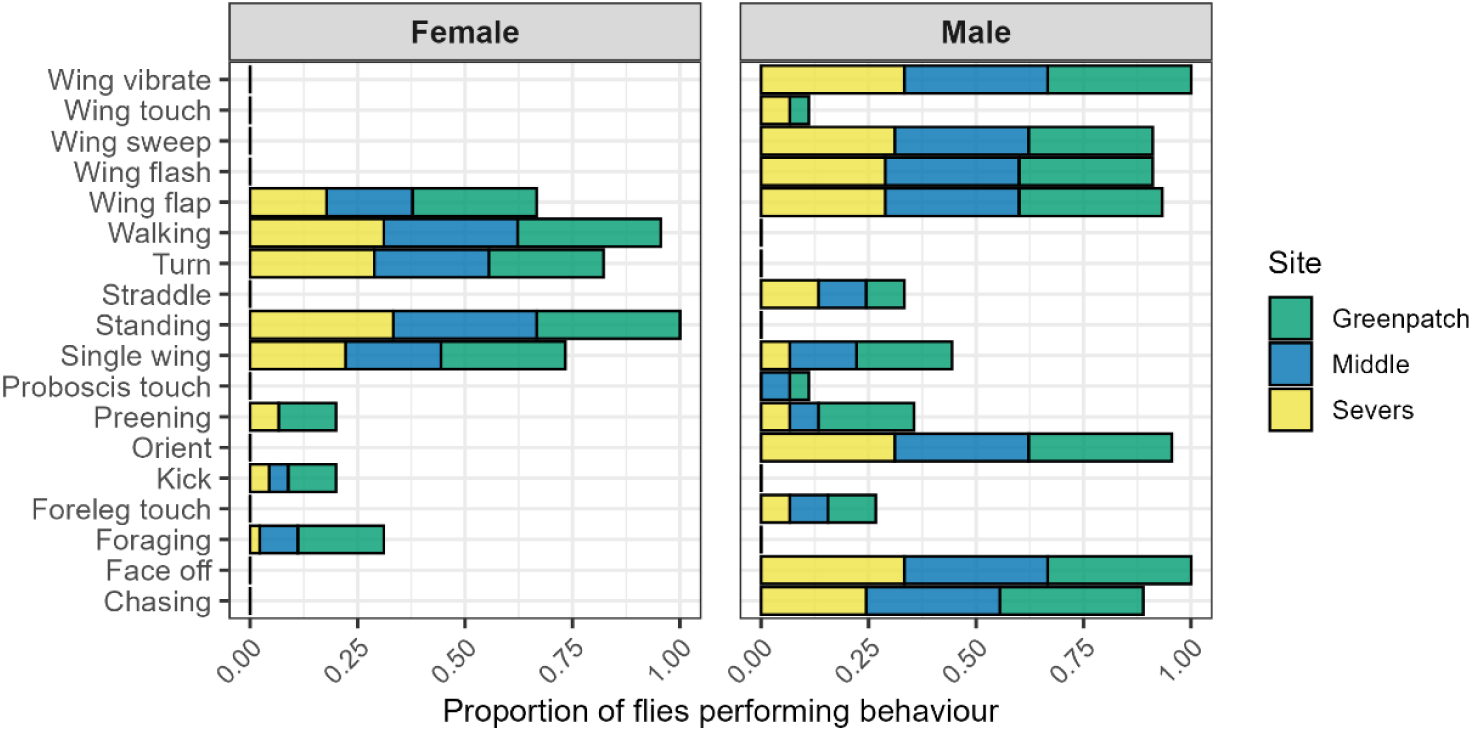
The proportion of male and female flies (out of all 45 males and females) exhibiting the 18 observed behaviours expressed during courtship. The relative contributions from each beach to the total proportions are represented in colour.

We found evidence that wing-sweep frequency differed among populations (Table 3; Figure 4A). This was due to a lower frequency of wing-sweep at Greenpatch than the other two populations and was driven by 3/15 individuals at Greenpatch Beach that only performed a single wing sweep during the display (Greenpatch 10, 12, and 14). However, wing sweep made up 50% of all behaviours expressed by Greenpatch 7 – so not all males at Greenpatch had low wing sweep frequencies. There were no significant differences in any other temporal patterns of the other main display components, including their duration, frequency, or interval (Table 3; Figure 4A), suggesting largely consistent temporal patterning among populations. Notably however, we identified high levels of variation in temporal patterning within- and among-populations.

**Figure 4.**
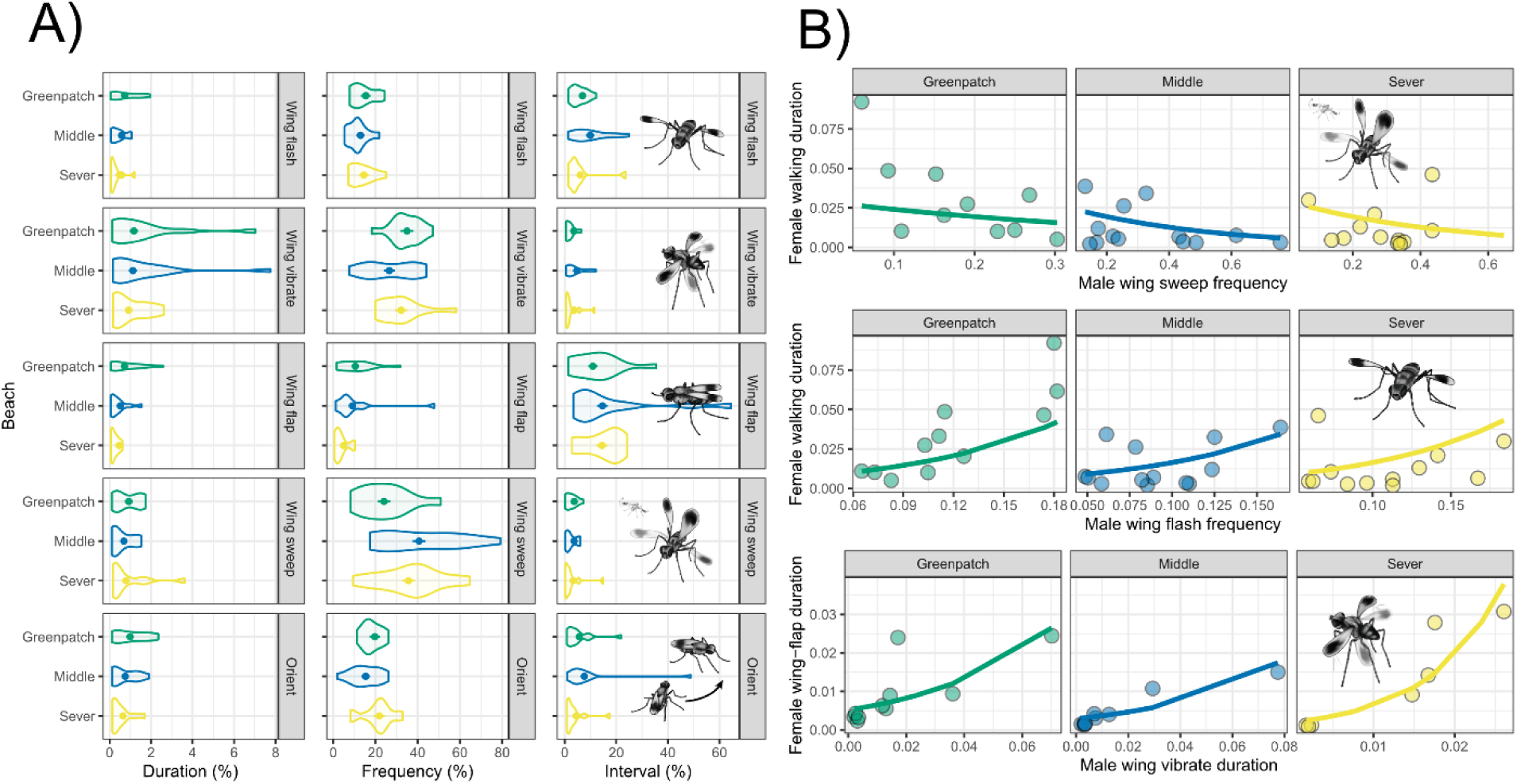
A) Predicted means and standard errors of the bout duration, frequency, and inter-bout interval of key male display components from the beta regression models (Table 3). B) Predicted fits of the beta regression models (Table 5) to the correlations between male display behaviours (wing sweep, wing flash, and wing vibrate) and female responses (walking and wing flap).

**Table 3.** Result of the analysis of variance (type III) for the effects of population (3 levels) and behaviour (5 levels) on mean proportional duration, frequency, and interval (Figure 4B). Bold numbers indicate significant values. To account for multiple comparisons, Bonferroni correction was applied (α = 0.01).

|  | Duration |  |  | Frequency |  |  | Interval |  |  |
| --- | --- | --- | --- | --- | --- | --- | --- | --- | --- |
| Full model | $\chi^2$ | df | p | $\chi^2$ | df | p | $\chi^2$ | df | p |
| Population | 2.44 | 2 | 0.295 | 1.69 | 2 | 0.429 | 2.69 | 2 | 0.261 |
| Behaviour | 7.59 | 4 | 0.108 | 87.24 | 4 | **<0.001** | 28.16 | 4 | **<0.001** |
| Population*Behaviour | 1.52 | 8 | 0.992 | 49.39 | 8 | **<0.001** | 6.56 | 8 | 0.584 |
| Wing flash |  |  |  |  |  |  |  |  |  |
| Population | 4.01 | 2 | 0.134 | 4.22 | 2 | 0.121 | 6.13 | 2 | 0.047 |
| Wing vibrate |  |  |  |  |  |  |  |  |  |
| Population | 0.97 | 2 | 0.616 | 7.15 | 2 | 0.028 | 3.2 | 2 | 0.202 |
| Wing flap |  |  |  |  |  |  |  |  |  |
| Population | 4.51 | 2 | 0.105 | 7.59 | 2 | 0.023 | 1.11 | 2 | 0.573 |
| Wing sweep |  |  |  |  |  |  |  |  |  |
| Population | 1.74 | 2 | 0.419 | 13.14 | 2 | **0.001** | 1.22 | 2 | 0.543 |
| Orient |  |  |  |  |  |  |  |  |  |
| Population | 3.44 | 2 | 0.179 | 3.24 | 2 | 0.198 | 7.24 | 2 | 0.0268 |

We found clear relationships between female preference proxies and male behaviours (Table 4, Table 5, Figure 4B). Wing sweep frequency and female walking bouts were negatively correlated (either females elicit male wing sweep when they stand, or male wing-sweep enhances female interest). Wing flash frequency and female walking bouts were positively correlated (either males use wing flashing to grab the attention of walking females, or male wing flash is a key component that causes female walking). Male wing vibrate bout duration and female wing flap bout duration were positively correlated (either male wing vibration elicits longer female wing flap bouts, or vice versa). Interestingly, in the one trial we observed to proceed to mating (Severs 1), the male spent the greatest proportion of his time (34%) in wing sweep, likewise the male in one of the longest trials before female rejection (576 seconds; Greenpatch 7) spent 50% of his time on wing sweep – thus, wing sweep is clearly an essential reflection of male courtship effort and subsequent female engagement in the display. However, we found no evidence that Greenpatch beach differed substantially in these patterns of female preference (Table 4), suggesting that the function of these traits and the female preferences are largely conserved across populations.

**Table 4.** Results of the analysis of variance (type III) for the effects of male display traits (wing sweep frequency, wing flash frequency, and proportion time spent wing vibrating) on female responses (proportion time spent walking and proportion time spent wing flapping) (Figure 4C). Bold numbers indicate significant values.

| Model, parameter | ANOVA (Type III) |  |  |
| --- | --- | --- | --- |
| | $\chi^2$ | df | p |
| **Female walking proportion ~ Male wing sweep frequency \* Population** |  |  |  |
| Male wing sweep frequency | 8.59 | 1 | **0.003** |
| Population | 11.46 | 2 | **0.003** |
| Male wing sweep frequency*Population | 5.54 | 2 | 0.063 |
| **Female walking proportion ~ Male wing flash frequency \* Population** |  |  |  |
| Male wing flash frequency | 20.15 | 1 | **<0.001** |
| Population | 0.18 | 2 | 0.912 |
| Male wing flash frequency*Population | 2.65 | 2 | 0.266 |
| **Female wing flap proportion ~ Male wing vibrate proportion \* Population** |  |  |  |
| Male wing vibrate proportion | 23.51 | 1 | **<0.001** |
| Population | 6.54 | 2 | **0.038** |
| Male wing vibrate proportion*Population | 24.27 | 2 | **<0.001** |

**Table 5.**
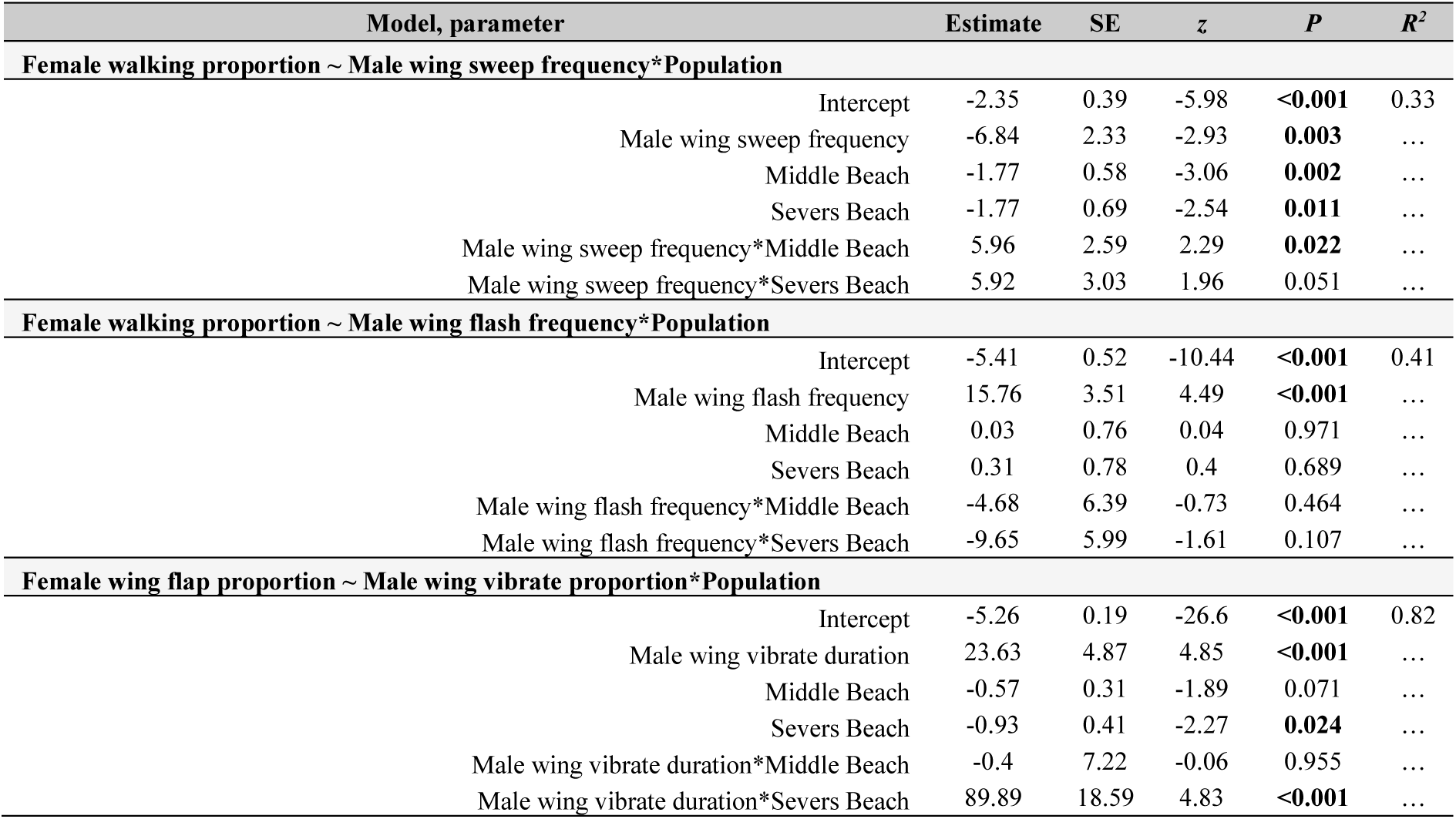
Results of separate beta regression models estimating the effect of male display traits (wing sweep frequency, wing flash frequency, and proportion time spent wing vibrating) on female responses (proportion time spent walking and proportion time spent wing flapping) (Figure 4C).

Finally, we found no evidence of divergence in the choreography or sequence of courtship displays among populations (Figure 5). Based on Markov analysis, the central well-connected nodes (behaviours) were the same across populations – with transitions occurring most frequently between the behaviours Face-off (6), Face-off-Wing-sweep (10), Orient-wing-vibrate (20), Face-off-Wing-vibrate (12), Orient-Wing-flash (17), Walking (34), and Wing-flap-Walking (38) (Table 6). Furthermore, no clear distinction in sequence across populations was detected based on Kullback-Leibler divergences, which were similar irrespective of the geographic distance of populations (mean KL divergences: Greenpatch x Middle: 0.179, Greenpatch x Severs: 0.184, Middle x Severs: 0.179) and we found no clear correlation between genetic distance and courtship distances (R^2^ = 0.074; Figure 5D).

**Figure 5.**
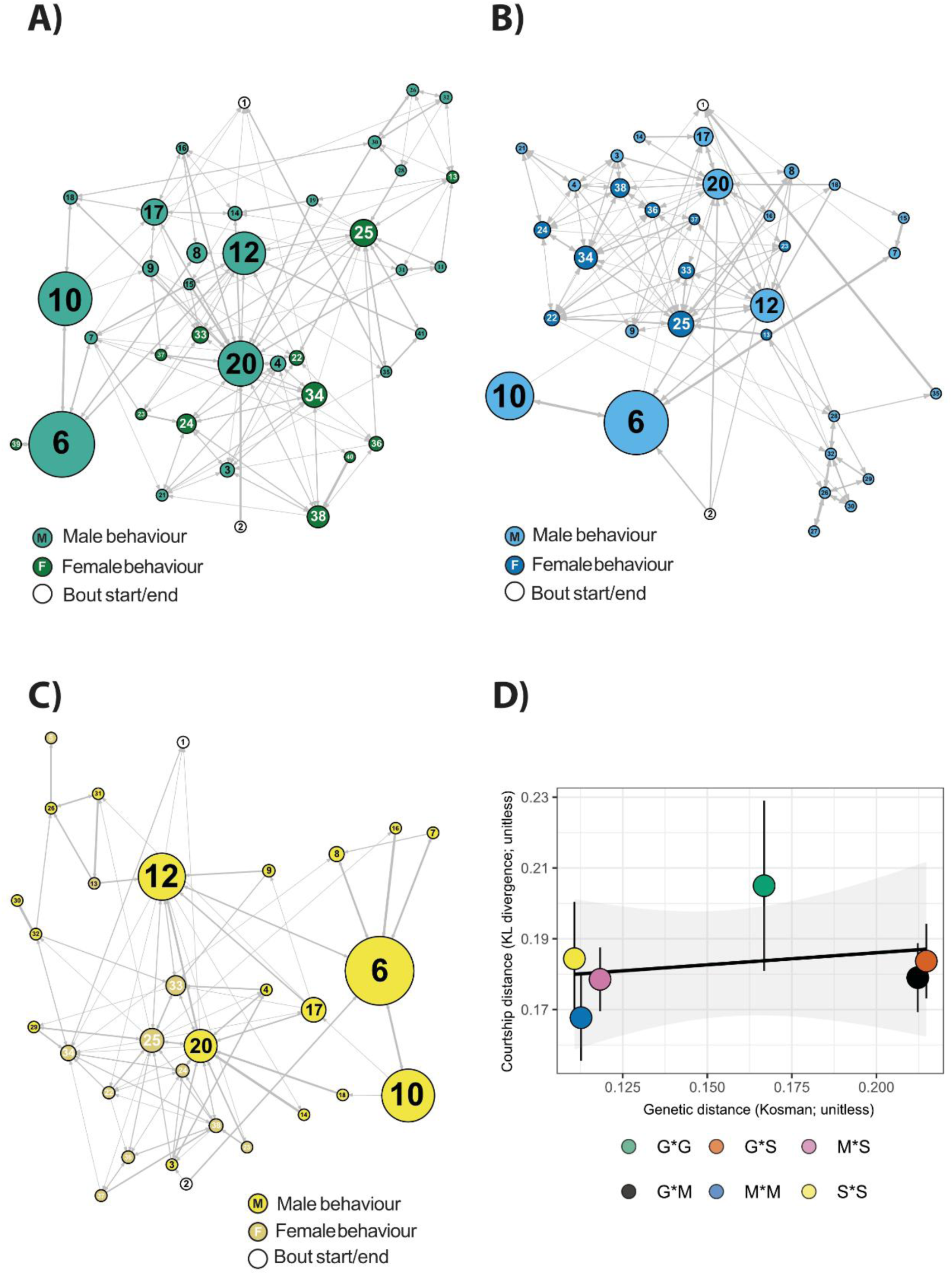
Sequential analysis of courting male-female pairs at A) Greenpatch Beach (G), B) Middle Beach (M), and C) Severs Beach (S). Circles with numbers represent discrete male and female behaviours (see Table 6). Circle size is scaled to the degree centrality of the behaviour (percentage of total courtship behaviours), lines represent significant transitions between behaviours, and line width is scaled to transitional probability (0 to 100%). Colour dictates whether the behaviour was performed by males or females; see legend of each plot. D) Mean inter-individual courtship distances ± SE (as calculated by Kullback-Leibler divergences of transition counts, 1 as pseudocount) against mean inter-individual genetic distances (individual genetic distances calculated by Kosman distance; Kosman & Leonard 2005) of each population pair. No clear relationship was observed between genetic distance and courtship distance (R^2^ = 0.074).

**Table 6.** List of behaviours in the sequential analysis (Figure 4A-4C).

| NUMBER | BEHAVIOUR |
| --- | --- |
| 1 | Bout-end |
| 2 | Bout-start |
| 3 | Chasing-(M) |
| 4 | Chasing-Wing-flap-(M) |
| 5 | Copulation-(F) |
| 6 | Face-off-(M) |
| 7 | Face-off-Single-wing-(M) |
| 8 | Face-off-Wing-flap-(M) |
| 9 | Face-off-Wing-flash-(M) |
| 10 | Face-off-Wing-sweep-(M) |
| 11 | Face-off-Wing-touch-(M) |
| 12 | Face-off-Wing-vibrate-(M) |
| 13 | Kick-(F) |
| 14 | Orient-(M) |
| 15 | Orient-Single-wing-(M) |
| 16 | Orient-Wing-flap-(M) |
| 17 | Orient-Wing-flash-(M) |
| 18 | Orient-Wing-sweep-(M) |
| 19 | Orient-Wing-touch-(M) |
| 20 | Orient-Wing-vibrate-(M) |
| 21 | Single-wing-(M) |
| 22 | Single-wing-Standing-(F) |
| 23 | Single-wing-Turn-(F) |
| 24 | Single-wing-Walking-(F) |
| 25 | Standing-(F) |
| 26 | Straddle-(M) |
| 27 | Straddle-Single-wing-(M) |
| 28 | Straddle-Wing-flap-(M) |
| 29 | Straddle-Wing-flash-(M) |
| 30 | Straddle-Wing-sweep-(M) |
| 31 | Straddle-Wing-touch-(M) |
| 32 | Straddle-Wing-vibrate-(M) |
| 33 | Turn-(F) |
| 34 | Walking-(F) |
| 35 | Wing-flap-(M) |
| 36 | Wing-flap-Standing-(F) |
| 37 | Wing-flap-Turn-(F) |
| 38 | Wing-flap-Walking-(F) |
| 39 | Wing-swing-Standing-(F) |
| 40 | Wing-swing-Walking-(F) |
| 41 | Wing-vibrate-(M) |

## DISCUSSION

There are perhaps almost as many unique courtship displays as there are animal species (Bastock 1967; Frith 1977; Arnold et al. 2017; Cannon 2023) yet we still know little about how such diversity arises in the face of constraining forces such as stabilising selection, pleiotropy, and structural interactions between courtship elements. Here, we investigated courtship divergence among three populations of the dancing dune fly *Apotropina ornatipennis* (Diptera: Chloropidae). Contrary to our expectation that courtship behaviour should diverge rapidly among allopatric populations, most measured aspects of courtship were consistent among populations despite clear genetic isolation by distance. This suggests that elaborate courtship displays can remain remarkably stable even in the face of genetic divergence.

Our results reinforce the notion that many components of courtship displays are conserved over short evolutionary timescales. We observed no additions or deletions of behavioural elements among populations (Figure 3). Likewise, except for differences in the frequency of ‘wing-sweep’, the timing of male behaviours (Figure 4A) and female responses (Figure 4B) were also consistent among populations, and the sequential structure of the display showed no correlation with genetic distance (Figure 5D). Similarly stable patterns of courtship have been observed in many other taxa, including in consistent expression of the songs of *Drosophila pseudoobscura* populations isolated for >75,000 years (Noor et al. 2000), the leg-waving displays of *Schizocosa crassipes* separated by over 1,000 km (Watts et al. 2019), and in the static calls of many species of hylid tree frogs (Gerhardt 1991). Such patterns of consistency also extend to macroevolutionary scales, such as in the phylogenetic conservation of the tail-straddling walk of *Plethodon* salamanders (Arnold et al. 2017), the bowing display of pigeons (Aves: Columbidae) (Goodwin 1966; Frith 1977), and in the neck displays of phrynosomatid lizards (Wiens 2000). Clearly then, courtship displays can be stable over vast evolutionary periods – but if so, when *does* the process of diversification begin, and how do courtship displays diversify so remarkably among species?

The only detectable divergence among our studied populations was in the frequency of ‘wing-sweep’ – with the northern population (Greenpatch beach) performing the behaviour less frequently than the two southern populations (Severs beach and Middle beach). Similar quantitative changes have been reported across the animal kingdom such as in the frequency of sigmoid displays of *Poecilia reticulata* (Luyten & Liley 1985), the duration of sine songs in *Drosophila teissieri* (Paillette et al. 1997), and the duration of the long chirp in *T. oceanicus* (Simmons et al. 2001). In these examples where courtship does diversify among populations, the changes are most often quantitative modifications of individual elements rather than wholescale rearrangements or deletions of display modules (Hebets et al. 2016; Arnold et al. 2017). Why? Possibly because, over short evolutionary timescales, it is unlikely that whole modules can be reorganised without impacting the functionality of the display (Arnold et al. 2017) – which could greatly reduce mating success and incur a substantial cost to fitness. Nevertheless, there is evidence that with sufficient ecological pressure, saltational jumps in courtship displays can occur among populations over very short timescales. In Hawaiian populations of *T. oceanicus* the entire courtship song was silenced over the span of 20 generations in response to exposure to a novel parasitoid (Zuk et al. 2006; Gallagher et al. 2024).

In the present study, the apparently low rates of courtship divergence may therefore simply be because of limited ecological variation among populations. Ecological variation is expected to be significant across latitudinal gradients – from variation in predation intensity, to the thermal environment and resource availability (Ketterson & Nolan 1976; Barnes 2002; Roslin et al. 2017; Freestone et al. 2021; Lush et al. 2024). Even slight differences in habitat characteristics like wind speed or frequency, sand colour, or vegetation types could lead to differences in male display trait optima among populations (as per sensory drive theory; Endler 1992; Cummings & Endler 2018). However, it is possible that beach environments are relatively homogeneous in ecological characteristics such as predation pressure, temperature, and resource availability – particularly over the short latitudinal scales in the present study (<1,000 km). In the context of background variation, which very likely varies among beaches, flies in the genus *Lispe* (Diptera: Muscidae) have shown the capacity to choose consistent display locations (performing their displays preferentially against dark backgrounds of seaweed wrack) which may buffer the signaller-receiver link even in the presence of substantial environmental variation (White et al. 2020) – and it is plausible that *A. ornatipennis* males elicit a similar strategy. The pace of divergence in courtship displays among the studied populations may thus be constrained by both the small scale of ecological differences across beaches and behavioural consistency in the microhabitat choice of displaying males.

Though environmental homogeneity alone cannot explain the lack of courtship divergence among populations – as theory predicts that mutation-order divergence (i.e., non-ecological diversification) should drive differentiation in sexual traits among populations even in the absence of ecological variation (Mendelson et al. 2014). It is therefore possible that courtship diversification is not stabilised by environmental homogeneity, but instead that there has simply been insufficient time for genetic drift or new mutations to occur in the alleles underpinning male display traits, or, that there are strong pleiotropic constraints (i.e., i.e., Yeh & True 2014; Ding et al. 2016) or structural constraints (i.e., Hebets et al. 2016) on male courtship traits slowing the pace of diversification. Equally, female preference landscapes could underpin the lack of mutation-order divergence. If genetic drift in female preferences is limited, or female preference landscapes are conserved among populations (as has been shown among some populations of *T. oceanicus* for the long chirp; Simmons et al. 2001), then even if new mutations underpinning male courtship traits do arise, they may be rapidly lost from the population.

Regarding the form of female preference in our populations, our results demonstrate high variance in the timing of male behaviours (Figure 4A) and show that females from all populations preferred extreme values of wing sweep frequency, wing flash frequency, and wing vibrate duration (Figure 4B). In line with this, the only male that successfully mated, and the male that held female attention for the longest duration (9.6 minutes) both had high wing sweep frequencies of 34% and 50% respectively. These patterns of high quantitative trait variance and female preference for extreme male values possibly indicate directional selection (Pomiankowski & Møller 1995) and may suggest that female preference functions in this system resemble a shallow linear or unimodal landscape that is conserved among populations – limiting the potential for alternative male trait peaks to be favoured across populations and constraining mutation order divergence.

Importantly, several caveats qualify these results. Female mating status was unknown, and copulations were rarely observed (mating was seen in only 1/45 trials). The majority of our observations therefore came from unsuccessful male-female pairs. It is possible that signals of divergence among populations in male traits and in female preferences may be detectable only when focusing on the subset of successful courtship displays that lead to mating. Although we did find evidence of divergence in wing-sweep frequency – even such differences must be interpreted with caution as the intensity and timing of male behaviours can be driven by differences in population density or other social factors that may vary among populations (Gerhardt 1991; Butterworth et al. 2019). Finally, while most measured aspects of courtship we measured did not detectably diverge among populations, female preferences very likely also depend on other courtship traits such as pheromones or acoustic cues (Arnold et al. 2017; Moran et al. 2020) which we did not measure. In *A. ornatipennis*, as in other chloropid flies, chemical and acoustic cues likely complement visual courtship (Kanmiya 1990; Yatsuk & Shestakov 2022), and a multimodal framework will be critical for future study. Likewise, high-speed videography and detailed kinematic analyses could reveal more subtle spatiotemporal features of male behaviours that are crucial components of female preference, and which diverge among populations.

Overall, our findings contribute to a growing recognition that many components of courtship are resistant to divergence under allopatry. This stability is consistent with uniform selection on choreography among populations, or genetic and structural constraints that limit divergence early in the process of speciation. In many situations therefore, courtship displays may be evolutionarily conserved, diversifying over much longer timescales or only under pronounced ecological variation among populations. Despite this, there are myriad examples of significant diversification in courtship displays among closely related species (Kusmierski et al. 1997; Ligon et al. 2018; Butterworth & Wallman 2021; Girard et al. 2021; Yukilevich 2021). To further elucidate the processes driving this remarkable diversification, comparative studies across populations and species using the framework we present here needed across a much wider range of taxa. Such work should involve observations of both sexes and integrate the full suite of courtship components - from differences in qualitative elements, to the timing of behaviours, and display sequence structure.

## Acknowledgements

NJB was granted permission to collect flies from Booderee National Park (Greenpatch Beach) as per the Environment Protection and Biodiversity Conservation Regulations 2000, Permit PA2021-00030. Flies collected from all other areas were permitted under the Biodiversity Conservation Act 2016, Permit SL101850.

NJB thanks Prof Phillip Byrne, A/Prof. Matthew Hall, Dr Chrissie Painting, and A/Prof Richard Peters for insightful discussions about the evolution of courtship displays. NJB thanks Kathryn Doty and Finlay Davidson for their assistance with fieldwork.

