## Supplementary material for "Courtship choreography is stabilised among genetically isolated populations"

**Table S1**. Pairwise Fst values.

|  | **SeversBeach** | **GreenpatchBeach** | **MiddleBeach** | **MerimbulaBeach** | **BearesBeach** | **HaywardsBeach** |
| --- | --- | --- | --- | --- | --- | --- |
| **SeversBeach** | *NA* | *NA* | *NA* | *NA* | *NA* | *NA* |
| **GreenpatchBeach** | 0.2666971 | *NA* | *NA* | *NA* | *NA* | *NA* |
| **MiddleBeach** | 0.0259084 | 0.2565752 | *NA* | *NA* | *NA* | *NA* |
| **MerimbulaBeach** | 0.0139292 | 0.277177 | 0.0133656 | *NA* | *NA* | *NA* |
| **BearesBeach** | 0.1603312 | 0.2834654 | 0.139685 | 0.1592649 | *NA* | *NA* |
| **HaywardsBeach** | 0.1682603 | 0.2810307 | 0.1448951 | 0.1675298 | 0.0207333 | *NA* |
| **WairoBeach** | 0.2429915 | 0.261379 | 0.2199907 | 0.2350984 | 0.1880884 | 0.175054 |
